# KIF20A cooperates with Myosin II to drive endosomal carriers fission and protein trafficking

**DOI:** 10.1101/2025.04.01.646333

**Authors:** Juliane Selot, Savinien Lasbareilles, Sabrina Colasse, Cédric Delevoye, Stéphanie Miserey, Joëlle Sobczak-Thépot, Anthi Karaiskou

**Affiliations:** Sorbonne Université, INSERM, UMR_S938, Centre de recherche Saint Antoine, CRSA, Paris, France; Université PSL, CNRS, UMR144, Institut Curie, Paris, France; Université Paris Cité, INSERM U1151-CNRS UMR 8523, Institut Necker Enfants Malades, INEM, Paris, France

## Abstract

Formation of membrane transport carrier relies on cytoskeleton coordination, however how microtubules and actin dynamics cooperate is still poorly understood. In this study, we reveal a novel mechanism by which a kinesin, KIF20A, regulates actomyosin and branched actin dynamics in coordination with Myosin II, facilitating the fission of transport intermediates at early endosomes. We demonstrate that KIF20A is required for maintaining endosomal homeostasis as its inhibition results in the enlargement of both early and late endosomes. Moreover, our findings demonstrate that KIF20A is required for the proper trafficking of Transferrin (Tf), and for ensuring the correct plasma membrane localization of β1 Integrin, a key protein that is critical for cell adhesion and migration. Collectively, these results highlight a new pivotal role of KIF20A as a key regulator of endosomal dynamics and function, contributing to metastatic potential.

## INTRODUCTION

During cancer progression, gene expression modifications, the loss of the original polarity and cytoskeleton reorganization are at the origin of the acquisition of a migratory phenotype, a prerequisite to metastatic events. Cell migration depends on the establishment of functional adhesions between cells and the extracellular matrix (ECM). These adhesions rely on Integrins, transmembrane proteins, which form a physical bridge between the ECM and the intracellular contractile acto-myosin fibers. In migrating cells, continuous reorganisation of the actin cytoskeleton and trafficking of membrane proteins, such as Integrins, promote the formation of new adhesions and protrusions on the cell surface (Caswell et al., 2009). The regulation of intracellular trafficking is therefore pivotal in tuning the final localization of Integrins, thereby influencing cell adhesion and motility dynamics.

The endo-lysosomal system is a complex network of interconnected organelles comprising early endosomes (EE), recycling endosomes (RE), late endosomes (LE), and lysosomes, responsible for determining whether internalized cargoes, such as solute molecules, ligands, and receptors, will be actively recycled or degraded (Maxfield and McGraw, 2004). Intracellular trafficking among endomembrane organelles and the plasma membrane relies on the formation of tubulo-vesicular transport carriers, which are eventually released, transported and delivered. These <delivery= carriers target diverse cell membranes either for material exchange or for the establishment and maintenance of molecular and functional identities of organelles. This process requires dynamic membrane remodelling events at multiple steps, including initiation and maintenance of membrane curvature to form a bud, extension of the bud to form a tubule, tubule neck constriction and finally fission to release the carrier. These mechanical modifications require the spatiotemporal coordination of the cytoskeletal network and associated proteins, which shape membranes through forces transmitted by molecular motors (McMahon and Gallop, 2005).

Actin filaments can actively deform membranes and are considered important contributors to endosomal membrane constriction and fission events (Simonetti and Cullen, 2019). Formation of branched actin filaments on endosomes, exerting a pushing force on the membrane, depends upon actin nucleators, such as the Arp2/3 complex, regulated by endosome-associated factors like the WASH complex (Derivery et al., 2009; Gautreau et al., 2022). Microtubules guide bud elongation and formation of tubule extensions in association with microtubule-based motors like dynein and kinesins (Delevoye et al., 2014). Myosin motor proteins can interact with actin filaments to generate the constriction forces required at the neck of the tubule (Ripoll et al., 2018). While actin polymerization is essential for tubule fission, excessive accumulation or stabilization can impair the process’s efficiency (Striepen and Voeltz, 2022). A tight coordination between actin, microtubules, and their motors and regulators, is needed to ensure the proper formation and release of transport intermediates (Delevoye et al., 2016).

Deregulations of motor proteins are frequently observed in cancers, disrupting cellular homeostasis. In particular, the overexpression of several kinesins has been strongly correlated with poor prognosis and survival outcome in patients with various cancers (Yu and Feng, 2010; Zhong et al., 2023). Among them, Kinesin family member 20A (KIF20A/Mklp2/Rab6KIFL) is upregulated in many cancer types and is repeatedly found within a group of genes associated with cancer progression and poor prognosis (Jin et al., 2023). KIF20A loss of function impairs cell division but also migration and invasion (Stangel et al., 2015; Duan et al., 2016).

KIF20A plays a crucial role during mitosis by facilitating the relocalization of the CPC (Chromosomal Passenger Complex) from the centromeres to the central spindle and the equatorial cortex (Serena et al., 2020). During cytokinesis, KIF20A promotes cleavage furrow progression via its ability to bind Myosin-II and actomyosin filaments (Kitagawa et al., 2013). In interphase, KIF20A has an additional function at Golgi fission hotspots, where it limits the diffusion of the Golgi-associated RAB6 GTPase on Golgi membranes and anchors RAB6 at microtubule nucleation sites with Myosin II to allow fission of post-Golgi carriers (Miserey-Lenkei et al., 2017). Both of the above studies describe a link between the microtubule and actin cytoskeletons to drive membrane constriction and fission events, mediated by the interplay between two molecular motors, KIF20A and Myosin II. While the essential role of KIF20A in cell proliferation can be explained through its well documented function in mitosis and cytokinesis, the molecular mechanism of KIF20A pro-migratory function remains elusive.

In this study we reveal a novel mechanism that regulates endosomal membrane fission, essential for β1-integrin protein trafficking at the plasma membrane. This mechanism relies on the coordination of the microtubule motor KIF20A and the actin motor Myosin II in the regulation of actomyosin contractility and branched actin dynamics at endosomal membranes. We demonstrate that KIF20A is essential for the recycling of cargos, such as integrins, which are crucial for the migratory phenotype and, consequently, the metastatic behaviour of cancer cells. The coordination of microtubule and actin cytoskeleton activities by molecular motors is a novel concept with significant implications for pathophysiology and could serve as a potential target for therapeutic strategies.

## RESULTS

### KIF20A activity is required for β1-integrin trafficking and cell adhesion and migration

We first investigated the role of KIF20A in cell adhesion and migration processes, using the Huh-7 cells as an *in vitro* model for hepatocellular carcinoma, with high KIF20A expression and motility potential (Gasnereau et al., 2012; Yau et al., 2013; Li and Wang, 2018). To directly assess the functional impact of KIF20A inhibition on cell adhesion, cells were exposed to Paprotrain, a highly selective pharmacological inhibitor of KIF20A (Tcherniuk et al., 2010) and were imaged during the spreading phase of adhesion. Time-lapse imaging of actin microfilaments was performed using SiR-actin, a live-cell actin probe. We observed that cells treated with Paprotrain appeared rounder, with a predominant actin belt, reminiscent of previous observations in bladder cancer cells (Mandal et al., 2019), with a smaller adhesion surface and delay in the establishment of lamellipodia (**Movie S1 and S2**). To quantify adhesion capacity, we performed real time adhesion assay (Dowling et al., 2014). As shown in **Fig. 1A**, PP-treated cells (red curve) exhibited a delay in adhesion, which became noticeable after 1 hour, compared to the control condition (blue curve). This effect was further quantified by calculating the area under the curve (**Fig. 1B**), confirming a reduction in adhesion potential upon PP treatment. Building on these results, we next investigated the impact of KIF20A inhibition on cell migration using a real-time migration assay. To define cell division-independent KIF20A- associated process, we treated cells with paprotrain and in parallel with mitomycin C to block cell proliferation, as prolonged KIF20A inhibition induces cytokinesis defects and increases cell size. Then, serum was used as a chemoattractant to induce cell migration. As shown in **Fig. 1C**, PP-treated cells (red curve) behaved like cells without chemoattractant (black curve), displaying a cell index four times lower than that of control cells (blue curve), indicating an inhibition of their migration capacity. This effect was further quantified by calculating the area under the curve **(Fig. 1D)**, confirming a strong reduction in migratory potential upon PP treatment. These data show that KIF20A is required for tumour cells adhesion and migration.

**Figure 1:**
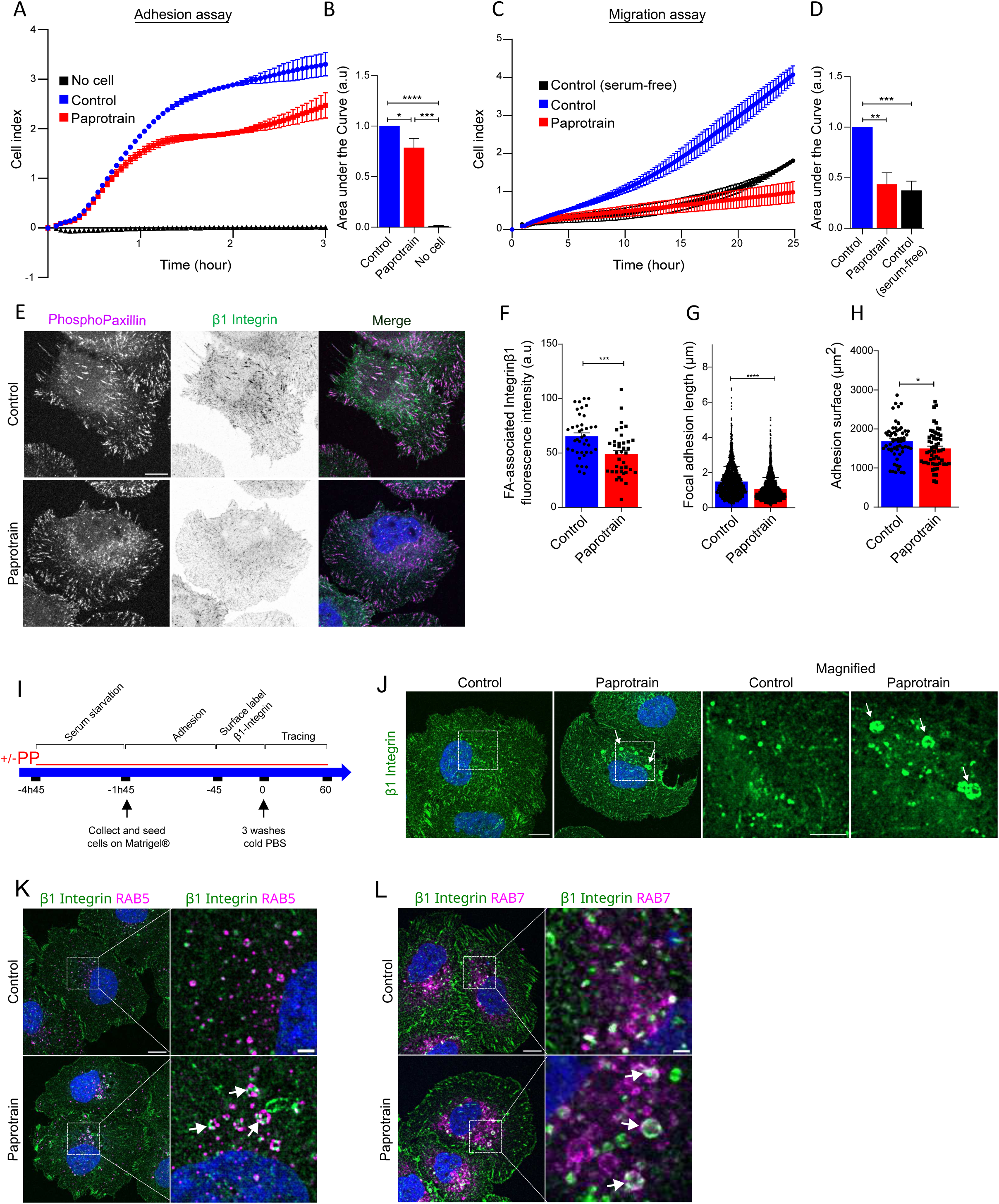
KIF20A activity is required for β1-Integrin trafficking, cell adhesion and migration. **(A)** Adhesion assay was performed using xCELLigence RTCA (Real-Time Cell Analysis). Huh-7 cells were treated with DMSO (control) or Paprotrain (25 µM) for 2hours. 2×10^4^ cells were seeded, and impedance was measured over 3h. The cell index reflects real-time adhesion based on impedance changes. The black curve represents well without cells. Data are shown as mean ± SD from 3/4 replicates in a representative experiment. **(B)** Quantification of the area under the curve from three independent experiments. Data are presented as mean ± SD, with statistical significance determined by an unpaired t-test. *p<0.05, ***p<0.001, ****p<0.0001. **(C)** Migration assay was performed using xCELLigence RTCA. Huh-7 cells, pre-treated with Mitomycin C (10 µg/mL) to block proliferation, were treated with DMSO (control) or Paprotrain (25 µM). 4×10⁴ cells were seeded, and impedance was measured over 24h with serum as a chemoattractant. The black curve represents spontaneous migration without a chemoattractant. Data are shown as mean ± SD from 3/4 replicates in a representative experiment. **(D)** Quantification of the area under the curve from three independent experiments. Data are presented as mean ± SD, with statistical significance determined by an unpaired t-test. **p<0.01, ***p<0.001. **(E)** Huh-7 cells were treated with DMSO (control) or Paprotrain (10 µM) for 2h, then detached and left to adhere onto Matrigel® (20µg/mL)-coated coverslips for 45 min. Cells were fixed and immunostained with antibodies against phospho-Paxillin and β1- Integrin. The images represent a summation projection over 3 slices, focusing on the plasma membrane in contact with the ECM. Scale bar = 10 µm. **(F)** Quantification of β1-Integrin at focal adhesion sites. Phospho-Paxillin was segmented to create a mask within which the raw intensity of β1-Integrin was measured (arbitrary unit). The histogram shows the mean ± SEM of two independent experiments. Statistical analysis was performed using an unpaired t-test ***p<0.001. **(G-H)** Quantification of the length of the focal adhesions **(G)** and the cell adhesion area **(H)** from Phospho-Paxillin signal shown in E. n= 60 cells per condition from three independent experiments, mean ± SEM. Statistical analysis was performed using a Mann- Whitney test **(G)** or an unpaired t-test **(H)**. *p<0.5, ****p<0.0001. **(I)** Experimental protocol of β1-Integrin antibody uptake assay. Huh-7 cells were serum-starved and treated with DMSO (control) or Paprotrain (25µM) for 2h. Cells were then detached and seeded onto Matrigel® (20 µg/mL) for 1 hour to assess adhesion. Cells were then incubated for 45 min at 4°C with β1- Integrin antibody conjugated to Alexa Fluor® 488. Medium was washed and cells were put back at 37°C to allow β1-Integrin endocytosis and recycling. Cells were fixed and analysed after 1h. **(J)** β1-Integrin tracking in control and paprotrain-treated Huh-7 cells, as described in **(H)**. Confocal microscopy images as the average intensity of 3 slices. Arrows indicated enlarged β1-Integrin vesicles. Scale bar = 10 µm, magnified inset scale bar = 5 µm. **(K-L)** Confocal images of Huh-7 cells treated as in **(I)** and immunostained for markers of early endosomes (RAB5, **K**) and late endosomes (Rab7, **L**). Scale bar = 10 µm. One slice is shown. Magnified views show the colocalization (in white arrows) between β1-Integrin and the markers (Scale bar = 2 µm).

During cell migration, cells must interact with and anchor to the ECM through adhesion sites known as focal adhesions. Among their key components, β1-integrin plays a crucial role in establishing functional focal adhesions by mediating cell-ECM interactions. To determine if KIF20A inhibition altered β1-integrin localization at the plasma membrane, which could explain the observed adhesion and migration defects, we quantified β1-integrin in mature focal adhesions, defined by the presence of phospho-paxillin, a key regulator of focal adhesion dynamics. During cell spreading, while the overall expression level of β1-integrin stayed constant (**Fig. S1, A and B**), KIF20A-inhibited cells exhibited a significant reduction (25 %) of the β1-integrin fluorescence intensity at the focal adhesion sites (**Fig. 1, E and F**; see methods for details). This was accompanied with shorter focal adhesions (**Fig. 1G**), and a reduction in the adhesion surface area (**Fig. 1H; Fig. S1, C and D**) compared to control cells, indicating a link between KIF20A activity and efficient β1-integrin participation in focal adhesions.

β1-integrin is submitted to continuous endocytosis and recycling at the plasma membrane which are essential for its proper distribution and function (De Franceschi et al., 2015). Any disruption in this balance can reduce β1-integrin expression at the cell surface. We thus monitored β1-integrin trafficking during cell adhesion, using an antibody uptake assay with a fluorescent anti-β1 integrin antibody (see M&M and **Fig. 1 I**). One hour after antibody uptake, labelled β1-integrin distributed as large intracellular vesicular structures in PP treated cells compared to control (**Fig. 1 J, arrows and S1E**). These structures were found positive for RAB5 and RAB7, well-characterized markers for early and late endosomes, respectively (**Fig. 1, K and L**). These results demonstrate that functional inhibition of KIF20A leads to aberrant intracellular accumulation of β1-integrin into large endosomal structures, suggesting a defect in its proper localization at the plasma membrane. This points to a disruption in the recycling process, where β1-integrin in the endosomes is likely in transit between endosomes and focal adhesions. This result indicates that KIF20A is involved in β1-integrin trafficking to the plasma membrane and localization at focal adhesion sites, leading to functional cell adhesion and migration.

### KIF20A localizes primarily to early endosomes

As HeLa cells reproduced the endosomal β1-integrin retention phenotype observed in Huh-7 cells upon KIF20A inhibition (**Fig. S1 F**), we used them as a model for intracellular trafficking studies. We first investigated KIF20A localization in HeLa cells by immuno-fluorescence microscopy (IFM) and focused on new endosomal positions that could be compatible with its implication in ꞵ1-integrin recycling. In interphasic cells, KIF20A has been reported to localize at the Golgi complex (Echard et al., 1998; Gasnereau et al., 2012; Miserey-Lenkei et al., 2017) (**Fig. S2A**). Interestingly, in PFA-fixed cells, we observed some additional cytoplasmic but non-Golgi-associated structures (**Fig 2. A-E**), that we characterized with well-characterized endosomal markers. We immunolabeled for both endogenous KIF20A and RAB5 and observed colocalization puncta (**Fig. 2A**), further confirmed with endogenous EEA1 staining of early endosomes (**Fig. 2B**). The corresponding line scan analysis demonstrates that the fluorescence intensity peaks are spatially superimposed, providing evidence for pixel-level colocalization (**Fig. 2, F and G, respectively**). To alternatively label EE, we transiently expressed in cells GFP-RAB5 and 2xFYVE-GFP. This resulted in an increase in the size of endosomes with an increased association of KIF20A (**Fig. 2, C and D**). Additionally, in order to examine endosomal compartments involved in the degradative pathway we labelled late endosomes/lysosomes with LAMP1-GFP (**Fig. 2, E and J**). Colocalization of KIF20A with this marker was limited, with occasional KIF20A spots on late endosomes, indicating a difference in KIF20A localisation between EE and LE. The Pearson correlation analysis between these markers and KIF20A reveals a significant differential distribution, with KIF20A preferentially localized to early endosomes (**Fig. 2K**).

**Figure 2:**
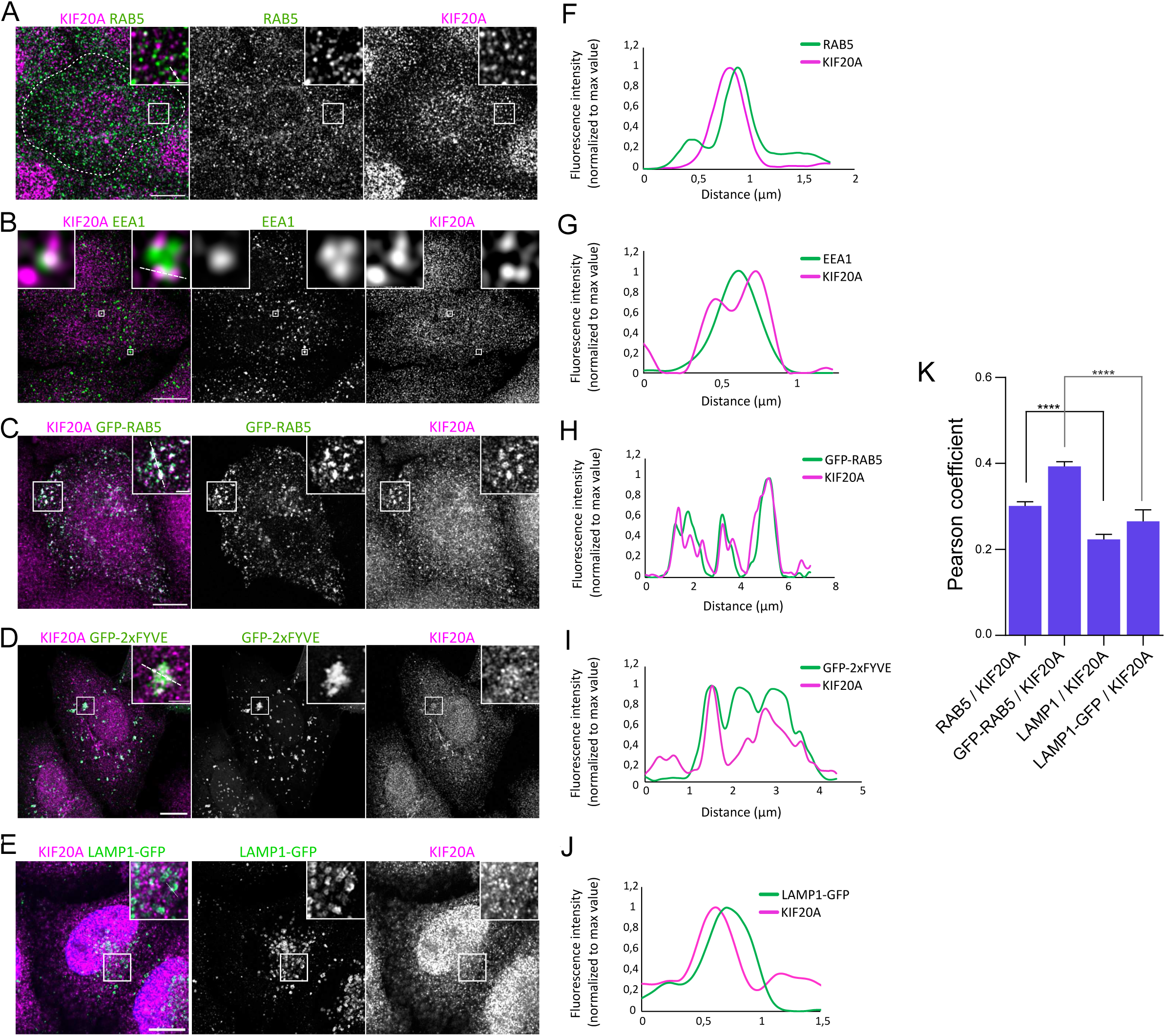
KIF20A localisation at endosomes. **(A-B)** HeLa cells were immunolabeled for endogenous KIF20A together with early endosomal markers: endogenous RAB5 **(A)**, and endogenous EEA1 **(B)**. Scale bar = 10 µm. Magnified images show partial colocalization between KIF20A and those markers (Scale bar = 2 µm). **(C-E**) HeLa cells were transfected with GFP-RAB5 **(C)**, 2x-FYVE-GFP **(D)** and LAMP1-GFP **(E)** expression vectors to visualize early endosomes. Scale bar = 10 µm. Magnified images show partial colocalization between KIF20A and those markers (Scale bar = 2 µm). **(F-J)** Line profiles of KIF20A and the corresponding markers fluorescence intensity (arbitrary unit, normalized to max value) along the white-dashed line showing the colocalization. **(K)** Pearson correlation coefficients between each co-staining, showing the degree of colocalization between KIF20A and the markers. Data are presented as mean ± SEM, with statistical significance determined by an unpaired t-test ****p<0.0001. n = 2-3 independent experiments, with a total of 40-100 cells analysed.

To validate the specificity of the KIF20A localization to endosomes, we performed a similar IFM experiment in cells following a siRNA-based silencing of endogenous KIF20A. This strategy resulted in the overall reduction of KIF20A signal both by western blotting and immunofluorescence (**Fig. S2B and C**), and the decrease of EEA1-associated KIF20A signal (**Fig. S2D**), further validating the specificity of KIF20A endosomal localization. Our data therefore reveal a novel endosomal localization of KIF20A suggesting a predominant role for KIF20A in early endosomal trafficking.

### KIF20A function is required for endosomal homeostasis

To investigate the primary function of KIF20A at endosomes, we examined changes in endosomal organization following KIF20A inhibition using Paprotrain. Inhibiting KIF20A activity for 2h caused a profound alteration of early and late endosomal structures (positive for RAB5, EEA1, GFP-RAB7 or LAMP1; Fig. 3A) characterized by an increase in size compared to controls, as shown by the distribution of EEA1+ and LAMP1+ vesicles, which exhibited a higher proportion of larger vesicles (>0.6 µm) (Fig. 3B). EEA1-positive endosomes showed a 2.1-fold increase in size (n ≥ 100 cells, 3 independent experiments), while LAMP1-positive endosomes exhibited a 2.4-fold increase in size (n = 40360 cells, 2 independent experiments) following KIF20A inhibition. This dynamic change in endosomal size was accompanied by a pronounced endosomal clustering in the perinuclear region. We followed size evolution during a 20-min kinetics, as growth rate analysis can provide insights into the underlying mechanism of the phenotype. This approach revealed that the increase in endosome size following KIF20A inhibition was rapid and progressive as reflected by the linear fit (**Fig. 3, C-E**). Furthermore, this phenotype was reversible after Paprotrain washout, with recovery occurring after 15 minutes for early endosomes (EEA1) and 45 minutes for late endosomes (LAMP1) (**Fig. 3, F and G**). In conclusion, inhibition of KIF20A activity causes a rapid and gradual increase in the size of endosomes, which is reversible when the activity is re-authorized and this with a distinct temporality between EE and LE, suggesting a primary defect in EEs with long term consequences in LE.

**Figure 3:**
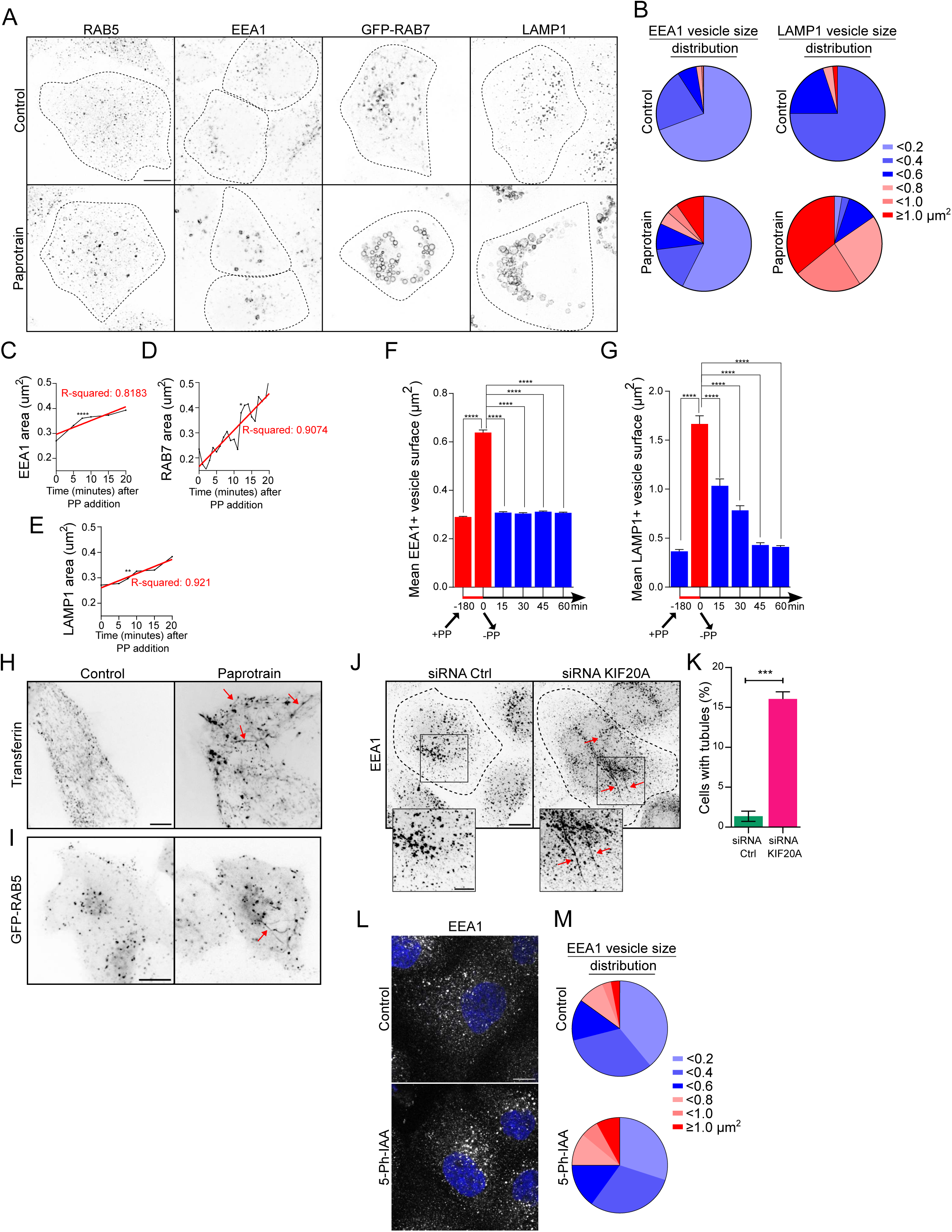
KIF20A activity is required for endosomal homeostasis. **(A)** HeLa cells treated with DMSO (control) or Paprotrain (25 µM) for 2h were either transfected with GFP-RAB7 and observed via live confocal microscopy or immunostained with antibodies against Rab5, EEA1 and LAMP1. The dotted lines represent the cell boundaries. The images represent a summation projection over 3 slices. Scale bar = 10 µm. **(B)** The distribution of EEA1 (across three independent experiments, n>100 cells), and LAMP1 vesicles (two independent experiments, 40-60 cells) according to their surface area was analysed in both conditions (DMSO and Paprotrain). The data show the frequency of vesicles across different size ranges. **(C-E)** Kinetics of endosomes enlargement. For EEA1 and LAMP1 vesicles, n>50 cells from each time point from 2 independent experiments. For GFP-RAB7, 5 cells were analysed from live imaging. Statistical analysis was performed using a Kruskal-Wallis test followed by Dunn’s multiple comparisons test. Statistical significance is indicated as follows: *p<0.05, **p<0.01, ***p<0.001, ****p<0.0001. The red line represents the linear regression of the data, and the R-squared value was calculated to assess the fit. **(F-G)** Time course of Paprotrain recovery. HeLa cells were treated for 2 h with Paprotrain (25 µM), washed, further cultured and fixed at various time points after paprotrain washout, and immunostained with antibody against EEA1 or LAMP1. Graphics show the surface area of EEA1-positive **(F)** and LAMP1-positive **(G)** vesicles. n = 30-66 cells per time points from two independent experiments. Data are presented as mean ± SEM. Statistical analysis was performed using a Mann-Whitney test ****p<0.0001. **(H)** HeLa cells were treated with DMSO (control) or Paprotrain (25 µM) for 15 minutes, followed by the addition of fluorescent Tf for 5 minutes. Cells were then imaged by time-lapse spinning disk confocal microscopy. The images represent the sum of 30 frames (1image/sec for 30sec). Long tubules containing Tf are observed upon KIF20A inhibition (red arrows). Scale bar = 10 µm. **(I)** GFP-Rab5 expressing HeLa cells were treated with DMSO (control) or Paprotrain (25 µM) for 20 minutes and then imaged by time-lapse spinning disk confocal microscopy (1image/2sec for 1min). A single frame was extracted from the movie. Long tubules are observed upon KIF20A inhibition (arrows). Scale bar = 10 µm. **(J)** HeLa cells were treated with control siRNA (Ctrl) or an siRNA targeting KIF20A, fixed and immunostained with an antibody against EEA1. The images represent a Z-projection over 10 confocal slices. Red arrows indicate tubulated endosomes. Scale bar = 10 µm, magnified inset scale bar = 5 µm. **(K)** Quantification of the percentage of cells exhibiting at least one visible tubule from experiment depicted in **(J)**. For each condition, 1500 cells were analysed across 3 independent experiments. Data are presented as mean ± SEM. Statistical analysis was performed using a paired t-test ***p<0.001. **(L-M)** MDCK KIF20A-GFP^AID^ were treated for 2h with DMSO (Control) or 5-Ph-IAA (auxin analog, 2 µM) to enable KIF20A degradation and immunostained with antibodies against EEA1 **(L)**. Scale bar = 10 µm. The distribution of vesicles according to their surface area was analysed in both conditions (DMSO and 5-Ph-IAA), across three independent experiments, n>200 cells. The data show the frequency of vesicles across different size ranges **(M)**.

A more detailed analysis of the dynamics of endosomal trafficking was conducted with live-cell imaging using fluorescently labelled transferrin (Tf) to track endosomes *via* an internalized cargo, while a separate set employed GFP-RAB5 to specifically label EE. HeLa cells treated with paprotrain (for 20 min) displayed long Tf-positive tubules as well as GFP-RAB5-positive tubules (**Fig. 3 H and I, and Movie S4 and S6**), absent in control cells (**Fig. 3 H and I, and Movie S3 and S5**), suggesting a defect in membrane fission. We next examined EE structure under conditions of siRNA-mediated KIF20A knock-down (as shown in **Fig S2**). Treatment with the siKIF20A resulted in the formation of long EEA1-positive tubules, observed in 15% of silenced cells compared to control cells (1%) rather than enlarged endosomes (**Fig. 3, J and K**), confirming the defect in membrane fission. The absence of the enlarged endosomes under these conditions could be due to the silencing kinetics which is slower compared to chemical inhibition but also to an incomplete protein depletion resulting in transient defects such as long tubules formation (**Fig. 3J and S2B, left panel**).

To further investigate the correlation between KIF20A inhibition strategy and the observed phenotype, we used a stable MDCK cell line expressing a degradable KIF20A carrying an AID tag together with a GFP tag (endogenous KIF20A with AID and GFP inserts). AID (Auxin Inducible Degron) tagged proteins become unstable thanks to ubiquitin-mediated proteolysis in the presence of auxin, enabling rapid, efficient and precise degradation of the protein of interest for functional studies (Zhang et al., 2015). As observed in **Fig. S3A and B**, KIF20A-GFP^AID^ showed partial co-distribution with EEA1-positive structures. As expected, upon a 2-hour incubation with 5-Ph-IAA (auxin analog), a global decrease of the KIF20A-GFP^AID^ fluorescence intensity was observed by IFM (**Fig. S3A and C**), paralleling a specific decrease of its intensity at EEA1-positives endosomes (**Fig S3D**). Importantly, this decrease within 2 hours of auxin treatment, was coupled with a change in the size distribution of EEA1 vesicles (**Fig 3, L and M**) with a higher proportion of large vesicles compared to control cells (25% of the vesicles were ≥ 0,6 µm^2^ in 5-Ph-IAA-treated cell, compared to 15% in control cell). These results using an alternative, precise and quick silencing of KIF20A, as opposed to a longer siRNA targeting approach, confirm that loss of function of KIF20A leads to enlarged endosomes, and reinforce the proposed role of KIF20A in endosomal homeostasis, potentially by facilitating the release of EE-derived transport intermediates through membrane fission.

### KIF20A regulates cargo sorting in early endosomes, influencing their trafficking and fate

The observed impact of KIF20A activity on early endosomal morphodynamics associated with a defect in β1-integrin membrane relocalisation, prompted us to explore endosomal sorting by conducting transferrin (Tf) and EGF recycling assays. We transiently exposed HeLa cells to fluorescently labelled Tf, that was then tracked and quantified based on image analysis (as described in **Fig. 4A**). Initial Tf uptake was not significantly altered in KIF20A-inhibited condition compared with control (**Fig. 4C**). However, a delay in Tf recycling was observed as early as 5 minutes post Tf endocytosis, with Tf retention in enlarged structures in the KIF20A- inhibited condition (**Fig. 4, B and D**). At each time point examined, KIF20A-inhibited cells contained a higher amount of intracellular Tf than control cells and the initial delay observed after 5 min of recycling did not significantly increase (**Fig. 4 D**) suggesting an impairment of fast recycling primarily.

**Figure 4:**
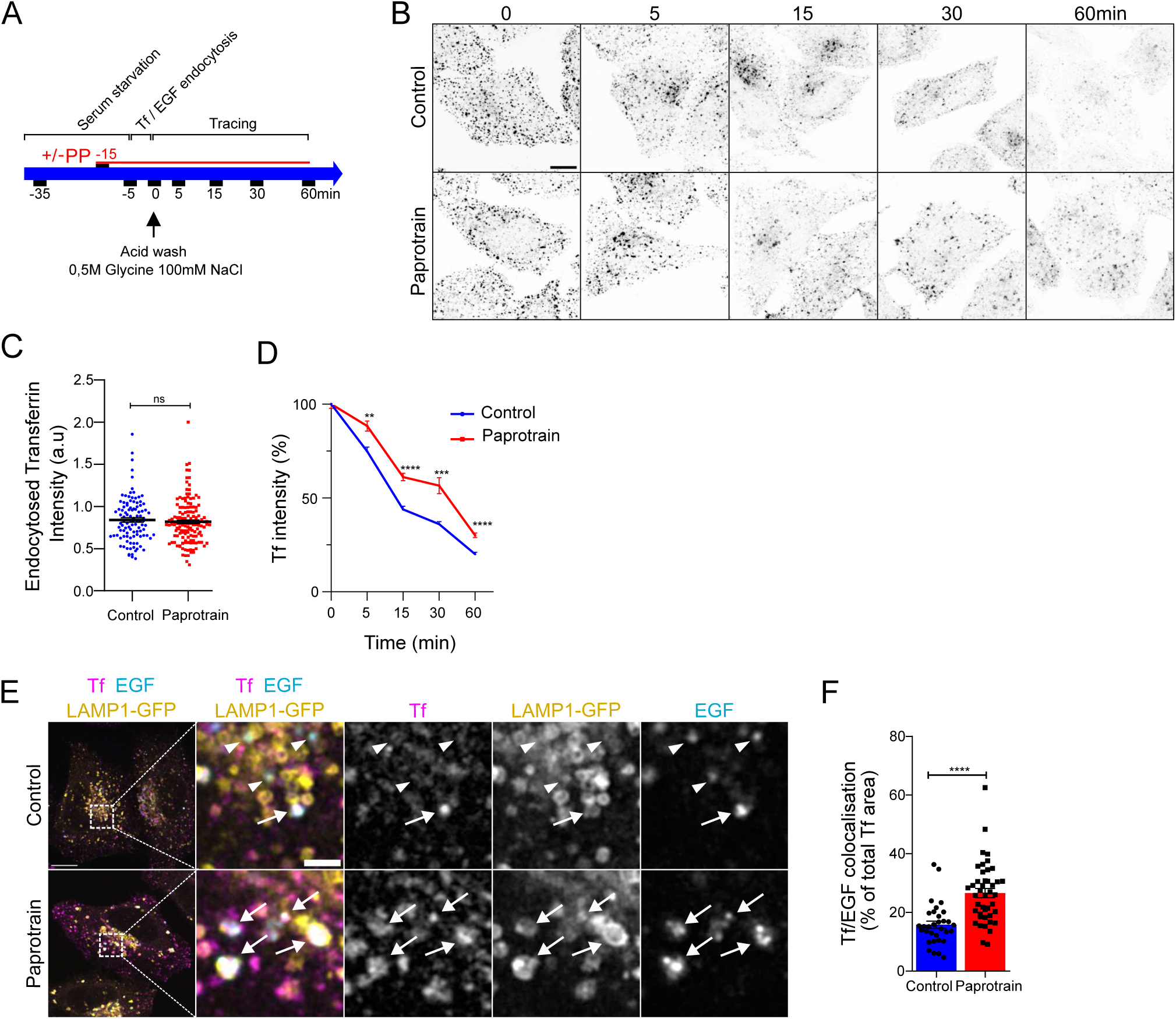
KIF20A regulates early endosomal cargo sorting. **(A-B)** Tf recycling protocol **(A)**. Briefly, HeLa cells were incubated with fluorescent Tf at 37°C for 5 min to allow endocytosis, followed by an acid wash at 4°C to remove membrane-bound Tf. After a shift to 37°C for the indicated times, cells were fixed and Tf recycling was observed from confocal microscopy images **(B)**. Scale bar = 10 µm. **(C)** Quantification of Tf endocytosis after the 5-minutes pulse (t0 of recycling). The graph shows the quantification from three independent experiments (n>100 cells). Data are represented as mean ± SEM. Statistical analysis was performed using a Mann-Whitney test to compare the conditions. ns = non significant. **(D)** Quantification of total Tf in cells treated with DMSO (control) or Paprotrain (25 µM) at various time points. The disappearance of fluorescent transferrin reflects the recycling of its receptor. The graph shows the quantification relative to time 0 from three independent experiments (n>100 cells). Data are represented as mean ± SEM. Statistical analysis was performed using a Mann-Whitney test to compare the conditions **p<0.01, ***p<0.001, ****p<0.0001. **(E)** HeLa cells were transfected with LAMP1-GFP and treated as in **(A)**, except that, in addition to Tf, EGF-A647 was added. Images show the cellular behaviour after 60 min of recycling (Scale bar = 10 µm). Magnified images show the colocalization between the 3 markers in white (Scale bar = 2 µm). Arrowheads indicate the colocalization between EGF and LAMP1, while arrows highlight the colocalization between EGF, Tf, and LAMP1. **(F)** Colocalization between Tf and EGF, measured as the % of Tf surface colocalized with EGF. n=32-43 cells from 2 independent experiments. Data are represented as mean ± SEM. Statistical analysis was performed using a Mann-Whitney test to compare the conditions ****p<0.0001.

We next explored the impact of KIF20A inhibition on both TfR and EGFR endocytic routes, internalized through clathrin-mediated endocytosis. After internalization, the two receptors segregate, with TfR being recycled back to the cell surface, while EGFR is directed to multivesicular bodies (MVB) and ultimately degraded in lysosomes (Futter et al., 1996). Cells were pulsed with fluorescent Tf and EGF ligands with the same protocol as above (Fig. 4A). As expected in control cells, while most of the internalized Tf was recycled to the plasma membrane, EGF was mainly localised in late endosomal/lysosomal LAMP1-positive compartments as evidenced by its colocalization with LAMP1 (Mander’s coefficient: 0,78 ± 0,07), and few colocalization spots positive for both Tf and EGF were observed (**Fig. 4 E**). In contrast, KIF20A inhibited cells exhibited a significant increase in co-localization between internalized Tf and EGF (**Fig. 4, E and F**, 60 min of chase) compared to control cells, suggesting that Tf is partially missorted to degradative compartments, reflecting a defect in cargo sorting. Together, our data support the conclusion that KIF20A is required for efficient cargo sorting and its inhibition leads to a delay in membrane protein trafficking.

### KIF20A and branched actin coordination for endosomal homeostasis

Cycles of polymerisation and depolymerisation of branched actin, through pro- and de- polymerising regulators are necessary for the fission of endosome-derived transport carriers required for cargo sorting and recycling to plasma membrane (Derivery et al., 2009; Simonetti and Cullen, 2019; Striepen and Voeltz, 2022).

We examined actin cytoskeleton organization in control and in KIF20A-inhibited cells by analysing F-actin and Cortactin status. Phalloidin labelling provides an overview of filamentous actin, whereas cortactin is a marker of branched actin structures. First, we observed that cortactin is localized to EE, together with KIF20A, as demonstrated by immunofluorescence staining of endogenous cortactin and KIF20A, on GFP-RAB5-labeled endosomes (**Fig. 5A**, white arrow highlights colocalization in the thresholded binary image). Interestingly, a marked increase in both F-actin and cortactin localizations to enlarged EE was observed following KIF20A inhibition by Paprotrain (**Fig. 5, B and C**). We analysed Cortactin intensity as a function of RAB5 vesicles size. Comparison of cortactin signal around large endosomes (> 0,7µm^2^) in control versus KIF20A-inhibited cells revealed a higher accumulation of Cortactin around large endosomes in paprotrain-treated cells (**Fig. 5D**). This finding suggests that KIF20A activity impacts branched actin polymerisation and supports the role of KIF20A in regulating actin dynamics at EE membranes.

**Figure 5:**
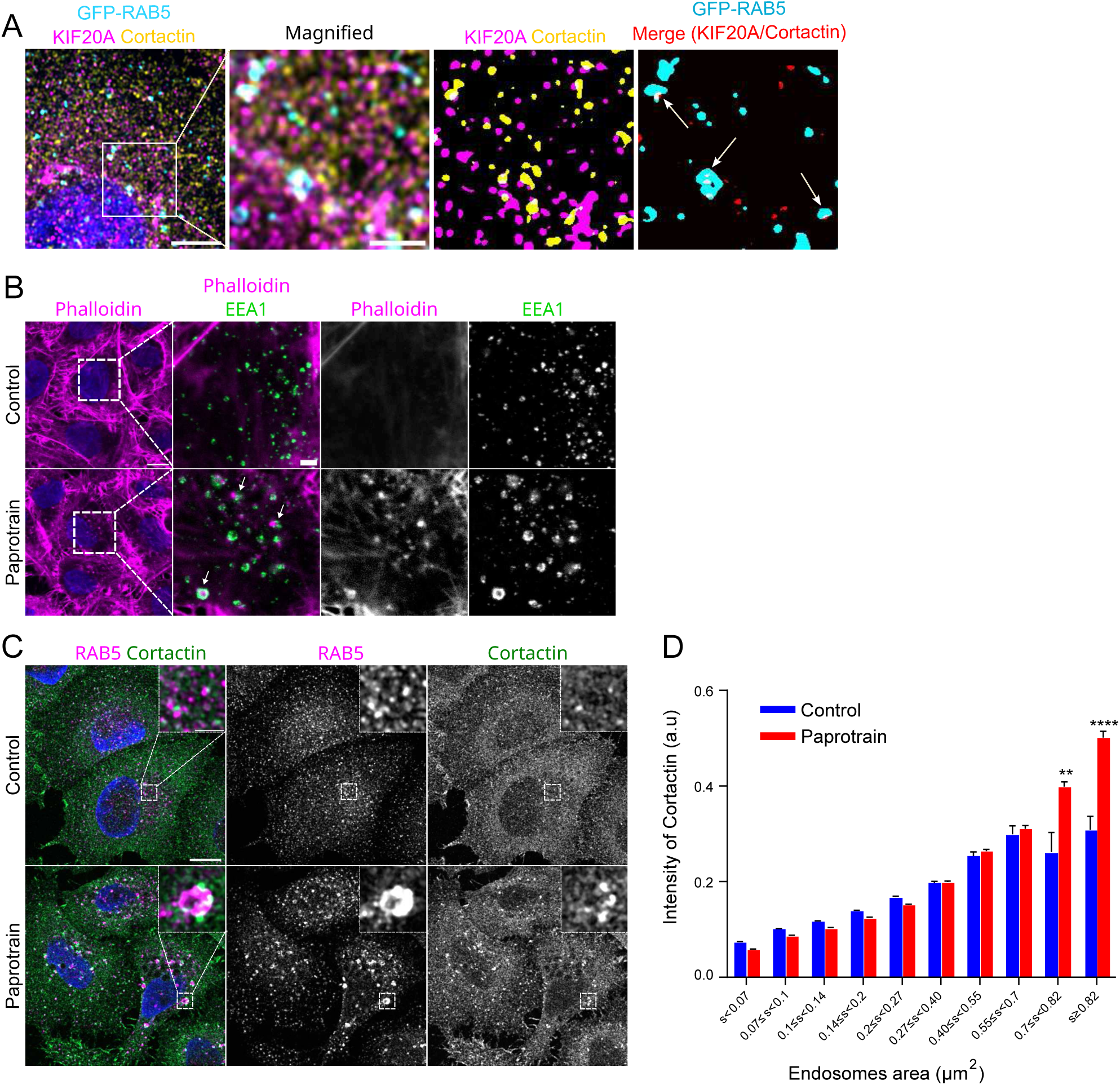
KIF20A activity modulates endosomal branched actin polymerisation. **(A)** HeLa cells were transfected with GFP-RAB5, detached, and left to re-adhere onto Matrigel-coated coverslips for 45min. Cells were then fixed and immunolabeled for endogenous KIF20A and Cortactin. KIF20A (in pink) and Cortactin (in yellow) were thresholded to create binary images, and the interaction mask was overlaid with the GFP-RAB5 mask. The white areas, indicated by the white arrows, indicate the colocalization of KIF20A and Cortactin at early endosomes. Scale bar = 10 µm, magnified inset scale bar = 5 µm. **(B)** HeLa cells were treated with DMSO (control) or Paprotrain (25 µM) for 2 hours, detached, and left to re-adhere onto Matrigel-coated coverslips for 45 min. Cells were then fixed and immunolabeled for endogenous EEA1 and actin was labelled with Phalloidin-A633. Scale bar = 10 µm, magnified inset scale bar = 2 µm. **(C-D)** HeLa cells were treated with DMSO (control) or Paprotrain (25 µM) for 2 hours, detached, and left to re-adhere onto Matrigel® (20 µg/mL) for 45 min, fixed and immunostained for endogenous RAB5 and Cortactin **(C)**. The RAB5 channel, created by summing three consecutive confocal slices, was segmented to identify individual vesicles. The cortactin signal was measured both within the vesicles and in a 0.3 µm surrounding field around each vesicle. Data were shown in the graph as a function of vesicle size **(D)**. mean ± SEM from three independent experiments, n>100 cells. Statistical analysis was performed using a Mann-Whitney test **p<0.01, ****p<0.0001.

### KIF20A and Myosin II cooperate at early endosomes

KIF20A is a direct partner of Myosin II at the equatorial cortex during cytokinesis (Kitagawa et al., 2013) and equally at Golgi-derived tubular carriers fission hotspots (Miserey-Lenkei et al., 2017). Recently, Myosin II assembly has been implicated in the generation of the mechanical forces required for fission events at endosomes in MDA-MB 231 cells (Kar et al., 2023). Therefore, KIF20A could cooperate with Myosin II to dynamically coordinate transport carriers formation/release at endosomes for cargos sorting and trafficking. We first reasoned that KIF20A and Myosin II should co-localize at early endosomes. By IFM, we indeed observed that endogenous KIF20A and Myosin Light Chain (MLC) partially co-distributed as dotty structures associated with GFP-2xFYVE labelled sorting endosomes (**Fig. 6 A**, white arrow highlights colocalization in the threshold binary image). Secondly, if KIF20A acts in coordination with Myosin II at endosomes, Myosin II activity inhibition would reproduce the endosomal phenotype induced upon KIF20A inhibition. MLC phosphorylation inhibition (thus actomyosin contractility inhibition) by a mix of ML74a Myosin Light Chain Kinase (MLCK) inhibitor4 and Y-276324A Rho kinase (ROCK) inhibitor, caused the enlargement of early and late endosomes, respectively marked by RAB5 and LAMP1 (**Fig. 6B**). This enlargement occurs as rapidly as KIF20A inhibition kinetics (**Fig. 6 C and D**), indicating a cooperation between these two motors in regulating endosomal size.

**Figure 6:**
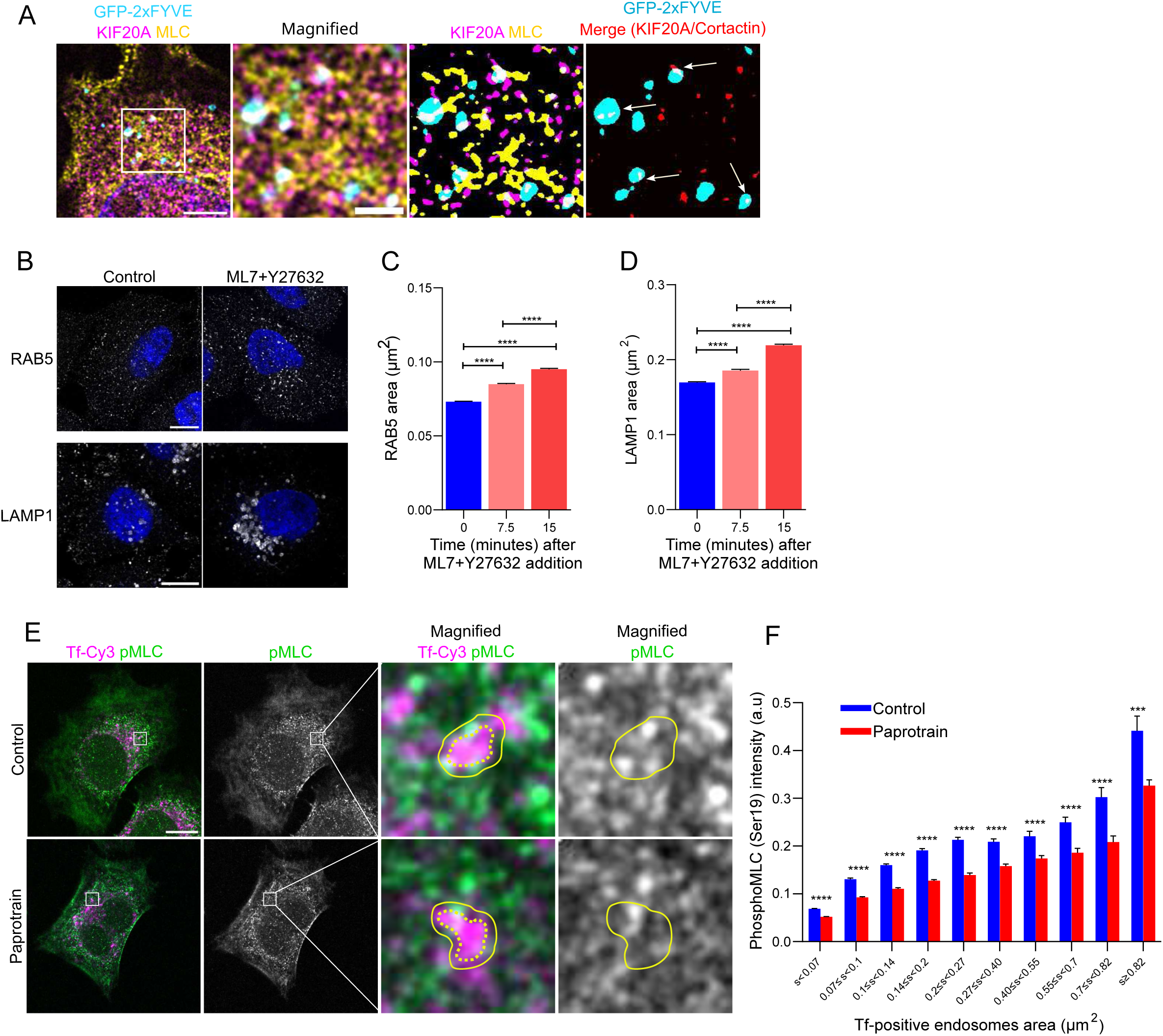
KIF20A and Myosin II cooperation at endosomes. **(A)** HeLa cells were transfected with GFP-2xFYVE to label EE, fixed and immunolabeled for endogenous KIF20A and MLC (Myosin Light chain). KIF20A (in pink) and MLC (in yellow) were thresholded to create binary images, and the interaction mask was overlaid with the GFP- 2xFYVE mask. The white areas, indicated by the white arrows, indicate the colocalization of KIF20A and MLC at EE. Scale bar = 10 µm, magnified inset scale bar = 5 µm. **(B)** HeLa cells were treated for 3hours with DMSO (control) or a mix of ML7 (30 µM) and Y27632 (10 µM), an MLCK inhibitor and a ROCK inhibitor, respectively. Cells were fixed and immunostained with antibody against RAB5 and LAMP1. Scale bar = 10 µm. **(C-D)** Kinetics of RAB5-positive **(C)** and LAMP1-positive **(D)** compartment enlargement after 7.5 and 15 minutes of ML7+Y27632 mix addition. Statistical analysis was performed using a Kruskal-Wallis test followed by Dunn’s multiple comparisons test. Statistical significance is indicated as follows: ****p<0.0001. **(E)** HeLa cells were treated for 2h with 25µM of Paprotrain, detach and re-adhere onto Matrigel® (20 µg/mL) for 45 min with Tf-Cy3, fixed and immunostained for PhosphoMLC (Ser19). The Tf channel from a single optical section localized at the center of the cell was segmented to identify individual vesicles (dashed line). Each vesicle was then expanded by 0.15 µm (straight line). **(F)** The pMLC signal was measured both inside the vesicles and within the 0.15 µm surrounding region and the data were shown in the graph as a function of vesicle size. mean ± SEM from three independent experiments, n >100 cells. Statistical analysis was performed using a Mann-Whitney test ***p<0.001, ****p<0.0001.

In order to further explore KIF20A and myosin II cooperation mechanisms, we assessed myosin II contractility in cells with inhibited KIF20A activity by measuring the overall level of phosphorylated MLC at Ser19 (pMLC) and more precisely its specific distribution at endosomes as a proxy of its local subcellular activity. Compared with control cells, Paprotrain- treated cells displayed a similar overall expression level of pMLC (data not shown). However, the fluorescence intensity of pMLC at endosomes loaded with Tf, specifically decreased in Paprotrain-treated cells, with a greater defect when endosomes are larger (**Fig. 6 E and F**). This result indicates that KIF20A activity is required for myosin II phosphorylation at early endosomes, likely promoting local actin-dependent contractility on endosomal membranes, which in turn facilitates transport carrier formation and fission (Kar et al., 2023).

## DISCUSSION

In this study, we present evidence that KIF20A, a microtubule motor protein, cooperates with Myosin II to regulate actomyosin contractility and branched actin polymerization at endosomes and thus intermediate transport fission. This new mechanism regulates cell membrane proteins availability, contributing to the acquisition of tumour cells motility phenotype, through β1- integrin recycling.

In this study, KIF20A knockdown strategies resulted in a combination of 3 major phenotypes: enlarged endosomes, endosome tubulation and defective cargo sorting suggesting an endosomal fission defect. The process of fission at endosomes is complex and not yet fully understood. Actin filaments drive membrane deformation and fission by generating pushing forces through branched filament assembly (Gautreau et al., 2022; Svitkina, 2018). Bud elongation and tubule formation rely on microtubular motors like dynein and kinesins. Constriction forces can also be generated by myosin proteins (Ripoll et al., 2018) and by recruitment of dynamin-like proteins following local depolymerisation of branched actin (Dhawan et al., 2022). Cellular phenotypes from different knockdown strategies of the above regulators include enlarged endosomes (Delevoye et al., 2016), endosome tubulation (Derivery et al., 2009; Freeman et al., 2014), or a combination of both (Fokin et al., 2021), mirroring those observed upon KIF20A inhibition. This places KIF20A as a novel regulator of fission dynamics. The reason for a tubulation versus enlargement phenotypes following KIF20A inhibition/knockdown is open to discussion but both are connected to defective cargo sorting and disrupted generation of transport intermediates. Cells expressing KIF20A at levels sufficient to prevent complete disruption of the endosomal network could display only transient defects, as long tubules formation. These long tubules generated over the long term can gradually reduce the size of the endosomes by consuming the excess membranes in the core of the EE.

We identified a novel localisation of KIF20A at endosomes, predominately on EE, distinct from its previously documented Golgi localisation (Miserey-Lenkei et al., 2017). While not specifically addressed here, localization and membrane recruitment of KIF20A on EE should be precisely regulated in time and space to appropriately control endosome-derived intracellular transport. For instance, KIF20A could be recruited to endosomal membranes through a direct interaction with its known partner, Myosin II, similar to the mechanism observed during anaphase (Kitagawa et al., 2013) or through direct interaction with phospholipids. Indeed, many motor proteins, including other kinesins (Hoepfner et al., 2005), rely on phosphoinositide interactions to achieve compartment-specific localization. Interestingly, the last 20 amino acids of KIF20A are enriched with conserved basic residues and referred to as the Lipid Association Motif (LAM), which could promote binding to various phosphoinositides, such as phosphatidylinositol 3-phosphate PI(3)P or PI(3,5)P_2_ (Fung et al., 2017; Tae et al., 2022) known to be respectively enriched at EE and LE. Further research into these mechanisms would be valuable to better understand the molecular processes involved in KIF20A’s recruitment to endosomal membranes.

A question that could be raised is the early versus late endosomal specificity of function of KIF20A. Our results indicate that this kinesin is primarily localized to early endosomes, and the formation of tubules was predominantly associated with these compartments. Furthermore, after an inhibitor washout, KIF20A activity is restored, and a reversion of endosomal enlargement is observed with distinct temporal dynamics between early and late endosomes, suggesting a primary defect in early endosomes, with long-term consequences for late endosomes. Furthermore, KIF20A inhibition results in a slowdown in the kinetics of Tf disappearance. This slowdown is observable from 5 minutes of recycling, suggesting an alteration mainly in the Rab4-dependent fast pathway. Consequently, the Tf clearance curves remain generally parallel throughout the kinetics, which does not indicate an additional defect that could affect, for example, the slow Rab11-dependent pathway. Although further analysis would be required, we propose that KIF20A exerts its effects primarily on EE and by regulating the fast recycling pathway.

How can this newly described endosomal KIF20A pool exert its function? We provide evidence that endosomal KIF20A co-localises with myosin II on EE. We further show that KIF20A activity is involved in endosomal actin reorganization, as its inhibition results in reduced actomyosin contractility associated with an accumulation of branched actin, both around the most enlarged endosomes. A counteracting crosstalk between actomyosin contractility and branched actin organisation has already been described in a biomimetic system (Sakamoto and Murrell, 2024) or during lamellipodia formation (Patel et al., 2021). In our context, the decrease in the contractile actomyosin network after inhibition of KIF20A could promote the organization of the actin network branched by Arp2/3, leading to an accumulation of the latter. Our data support a new mechanism of cooperation of two molecular motors (KIF20A-Myosin II) that could generate and transmit forces modulating branched actin dynamics at the level of endosomes. This process quickly impacts the fission of transport intermediates, thereby maintaining endosomal homeostasis, regulating cargo sorting and transport, and ultimately impacting cell adhesion and migration.

It has been shown that KIF20A participates in the fission of Rab6-positive vesicles of the Golgi apparatus in association with myosin II (Miserey-Lenkei et al., 2017), and this in combination with our results suggest that KIF20A may act as a general regulator of actin dynamics for tubular transporter fission. A similar mechanism could be present in different organelles, with distinct dynamics highlighting regulatory processes that are not fully understood. It would be relevant to examine the involvement of KIF20A in other organelles undergoing fission events, such as mitochondria, especially since Myosin II has already been associated with the regulation of mitochondrial fission (Korobova et al., 2014).

Finally, we show that KIF20A motor activity inhibition leads to impaired ꞵ1-integrin trafficking, cell adhesion and migration, highlighting the importance of KIF20A in regulating key processes involved in cell motility and metastatic potential. KIF20A is overexpressed in a variety of malignant tumours and is associated with poor patient prognosis and metastatic phenotypes. Inhibiting KIF20A expression has been shown to down-regulate proliferation and invasion of breast or pancreatic cancer cells (Stangel et al., 2015; Nakamura et al., 2020). For the first time here, we provide a mechanistic explanation of the involvement of KIF20A in dynamic cell motility, a property required for metastatic cells. We propose that overexpression of KIF20A in cancer cells acquiring migratory capacity can support β1 integrin recycling processes. We show that trafficking of other membrane proteins, such as TfR, also depends on KIF20A activity. It would be interesting to study trafficking dynamics of the second family of key metastatic membrane proteins, the metalloproteases.

Our study reveals a novel mechanism by which, KIF20A, a microtubule motor protein in coordination with an actin motor, Myosin II, influences actin network dynamics to control tubular carrier fission and membrane protein recycling, providing new insights into the coordination of cytoskeletal and membrane remodelling processes.

## EXPERIMENTAL PROCEDURES

### Cell lines

HeLa Kyoto cells were a kind gift from Olivier Gavet (Gustave Roussy, France) and MDCK KIF20A-GFP-AID cell line was from Benedicte Delaval and Benjamin Vitre (CRBM, Montpellier, France). Cells were grown at 37°C in 5% CO_2_ in DMEM GlutaMAX^TM^ (Gibco) supplemented with 10 % fetal bovine serum (FBS) and 100 µg/mL penicillin/streptomycin. Human hepatoma cell line Huh-7 was maintained in minimal essential medium containing Earle’s salt, 1% nonessential amino acids, 1 mmol/L sodium pyruvate, 10 % FBS and 100 µg/mL penicillin/streptomycin.

### Transfection

Approximately 3×10^5^ cells were transfected with 0,25 µg of plasmid DNA using XtremeGENE HP (Roche) according to the protocol provided by the manufacturer (reverse transfection). The cells were analysed 24h post transfection. For the silencing experiments, 1,6×10^5^ cells were transfected with 20 nM of KIF20A siRNA using Lipofectamine^TM^ RNAiMAX (ThermoFisher) twice at 24h interval and analysed 24h after the last round of transfection.

### Plasmids DNAs and siRNAs

The plasmids encoding the following fusion proteins were used: GFP-RAB5, GFP-RAB7, LAMP1-GFP, 2x-FYVE-GFP (Institut Curie, France). mRFP-RAB7 was a gift from Ari Helenius (Addgene plasmid # 14436).

The sequence of the siRNA used in this study is the following: human KIF20A: (3’- GGAACATAGTCTTCAGGTA-5’) synthesized by Dharmacon Research, Inc.

### Antibodies

The following antibodies were used:

Rabbit polyclonal anti-KIF20A (1:2000 IF), a generous gift from A. Echard (Institut Pasteur), Rabbit anti-GFP (Invitrogen, #A-11122; 1:200 IF), mouse anti-cortactin (Sigma-Aldrich, #05- 180-I; 1:500 IF), mouse anti-EEA1 (BD Biosciences, #610457; 1:500 IF), mouse anti-GM130 (BD Biosciences, #610823; 1:500 IF), mouse anti-β1-Integrin (#MAB2253, Invitrogen, 1:300 IF). Mouse anti-Myosin Light Chain was from Abcam (#ab89594; 1:300 IF). Rabbit anti-Filamin was from Genetex (#GTX101206; 1:300 IF).The following antibody were obtained from Cell Signaling: rabbit and mouse anti Rab5 (#3546 and #46449, respectively; 1:500 IF), rabbit anti-Rab7 (#9376; 1:100 IF), rabbit and mouse anti-LAMP1 (#9091 and #15665, respectively; 1:500 IF), rabbit anti-PhosphoPaxillin (#69363, 1:300 IF), rabbit anti-Phospho-Myosin Light Chain (Ser19) (#3675; 1:300 IF). Mouse anti-Lamin A/C (sc-376248, 1:500 WB), anti-GAPDH-HRP (sc-47727; 1:10000 WB) and anti-Actin-HRP (#sc-1615; 1:1000WB) antibodies were obtained from Santa Cruz. Mouse anti-EGFR was from ProteinTech (#66455; 1:10000 WB). The β1-integrin antibody used for Western Blot (1:500) was a generous gift from Corinne Albigès-Rizo (Institut pour l’Avancée des Biosciences, Université Grenoble Alpes). The secondary antibody used for IFM were obtained from Invitrogen: anti-mouse IgG A568 (#A11004), anti-rabbit IgG A568 (#A11011), anti-mouse IgG A488 (#A11001), anti-rabbit IgG A488 (#A11008), anti-mouse IgG A647 (#A21235), anti-rabbit IgG A647 (#A21244) and used at 1:500 dilution.

For Integrin trafficking, Alexa Fluor® 488 anti-human CD29 antibody (# 303016, 1:100 IF) was obtained from BioLegend.

### Chemicals and recombinant proteins

The cell permeable acrylonitrile inhibitor of KIF20A (a.k.a., Paprotrain for PAssenger PROteins TRAnsport Inhibitor, Tocris) was solubilized in DMSO as a 100 mM stock solution protected from light due to its photosensitivity, and stored at -20°C. To inhibit KIF20A, cells were cultured in the presence of 10 or 25 µM inhibitor. Incubation times ranged from 5 minutes to 3 hours as mentioned in figure legends. The MLCK inhibitor ML7 was purchased from Sigma (#I2764) and the ROCK inhibitor Y-27632 was purchased from Clinisciences (HY- 10583). 30 µM ML7 and 10 µM Y-27632 was added to the cell medium at 37°C under 5% CO_2_. Time incubation are mentioned in the legend.

To induce degradation of endogenously tagged KIF20A-AID-GFP, 2 µM of 5-Ph-IAA (MedChemExpress # HY-134653) was added directly to the culture medium for 2h. Human Transferrin-A488 and Transferrin-Cy3^TM^ were obtained from Jackson IR (#009-540- 050 and #009-160-050 respectively). Human Alexa Fluor™ 647-EGF complex and Phalloïdine-A633 were obtained from ThermoFisher (# E35351 and # A22284 respectively).

### Cellular Lysis and Western blotting

For preparation of cellular lysates, cells were lysed at 4°C in RIPA buffer (Bio Basic Inc. Cat# RB4475) supplemented with 1X Halt™ Protease and Phosphatase Inhibitor Single-Use Cocktail (ThermoFisher Cat#78442), 1X EDTA, 0,5% sodium deoxycholate for 5 min, harvested, and lysed for 5 more minutes. Extracts were then centrifuged at 13.000 rpm for 10 min and supernatants were retained. To isolate nuclear and cytosolic fraction, we used the Subcellular Protein Fractionation Kit for Cultured Cells (ThermoFisher Cat# 78840). Protein estimation was performed with BCA protein quantification assay kit (ThermoFisher Cat #23225). The lysates obtained were boiled with 4xLaëmmli buffer (8% SDS, 0,2M Tris-HCl pH 6.8, Glycerol 40%, 0,4M DTT, bromophenol blue) for 5 min at 95°C and then resolved by SDS- PAGE. The resolved proteins were transferred to nitrocellulose membrane and then incubated overnight with primary antibodies. The blots were then probed with HRP tagged secondary antibodies (Cell signaling, anti-mouse IgG #7076S, anti-rabbit IgG #7074S; 1:10000 WB) and chemiluminescence signals were captured by a Chemidoc system (Bio-Rad, USA). Western-blot quantification was performed by measuring the adjusted intensity using ImageLab (Bio-Rad, USA).

### Endocytosis and recycling assay

#### β1 Integrin trafficking

Cells were serum-starved and treated with DMSO (control) or Paprotrain (25 µM) for 3h. Cells were then detached and re-adhered onto Matrigel® (20 µg/mL) for 1 hour to assess adhesion. Cells were then incubated for 45 min at 4°C with β1-Integrin antibody conjugated to Alexa Fluor® 488. Cells were washed and either fixed (t0) or further cultured at 37°C with fresh medium (+/- PP) and fixed after 1h.

#### Transferrin assay

HeLa cells grown on glass coverslips were serum starved for 30 min at 37°C in DMEM supplemented with 1% BSA and then incubated with DMSO (control) or Paprotrain (25 µM) for 10 min. Cells were further incubated in DMEM/BSA medium with Tf-Cy3 or Tf-A488 at a final concentration of 5 µg/mL (5 min, 37°C, 5% CO_2_) to allow Tf endocytosis. Then endocytosis was stopped and membrane bound Tf was removed by acid-wash (50 mM Glycine, 100 mM NaCl, pH 3.0, 10 sec) and washed 3 times in cold PBS (ThermoFisher). Cells are fixed to examine endocytosis or incubated in medium supplemented with serum to follow Tf recycling at different time points. Paprotrain or DMSO vehicle were kept for the whole experiment. Alternatively, cells were pulse labelled with Transferrin and EGF-Alexa647 (2 µg/mL).

#### Migration assay using Real Time Cell Analysis (RTCA)

To irreversibly arrest cell proliferation, Huh-7 cells were treated with Mitomycin (10 µg/mL, 3 h), then washed twice with PBS, serum starved and pre-treated with DMSO or Paprotrain (25 µM, 3 h). Cells were then seeded in DP plates (Agilent) and cell migration was monitored in real time as described by the manufacturer. Briefly, 3×10^4^ cells were added in the upper chamber coated with a low concentration of Matrigel (20 µg/mL) to facilitate Paprotrain-treated cell adhesion. Medium containing 10% serum as chemoattractant was added in the lower chamber. Serum-free medium was used as negative control to measure spontaneous migration. Each condition was performed in triplicate. The migration rate was assessed by measuring electrical impedance due to cell migration and adhesion on gold electrodes underside of the membrane which separated the two chambers. Original datasets expressed in arbitrary units (cell index) were exported to MS Excel and the raw data were processed without CI- normalization.

#### Immunofluorescence and confocal microscopy

HeLa cells were seeded on matrigel-coated coverslips one day before the experiment unless otherwise specified. Cells were fixed either in warm 4% para-formaldehyde for 15 min at RT or in methanol for 5 min at -20°C. Immediately following fixation, cells were either permeabilized for 5 min with 0,2% Triton X-100 in PBS followed by 30-min saturation (3% BSA in PBS) or permeabilised and saturated for 30 min with 0.05% saponin, 3% BSA in PBS. Cells were then incubated in PBS/BSA or PBS/BSA/saponin with the primary antibody overnight at 4°C. Coverslips are then washed three times in PBS or PBS/saponin with gentle shaking and incubated with the corresponding secondary antibody for 45 minutes. Cells were again washed three times in PBS and incubated for 10 minutes in PBS supplemented with 1 µg/mL Hoechst33342 (ThermoFisher) for nucleus visualization. Finally, coverslips were mounted in Prolong Antifade Reagent (ThermoFisher). Phalloidin-A633 is used for the visualization of the F-actin and added for 30 min together with Hoechst 33342.

Fluorescence cells images were obtained with a LSM 980 Zeiss confocal microscope with the Airyscan 2 technology. We used an 63x oil immersion NA 1,4 objective, the 405, 491, 561, 639 excitation laser lines. Image acquisition was performed with an optimal step size of 0,14 µm. Airyscan image processing from raw data was performed with ZenBlue software. For some experiments (endosomes kinetics), images were acquired using a Fluoview 3000 confocal microscope (Evident). A 60X UPlan Apo NA 1.42 oil immersion objective was used, with the 405, 488, 561 excitation laser lines. Image acquisition was performed with an optimal step size of 0.4 µm, as determined by the software.

Quantification of (1) HeLa cells exhibiting tubular structures after KIF20A silencing and (2) the surface of Huh-7 cell adhesion after treatment with paprotrain was performed on galleries comprising over 500 cells, acquired on an inverted IX83 microscope (Olympus) with 60XS2 Silicone WD 0,3mm objective, driven by Cell Sens dimension V1.14 software.

#### Time-lapse imaging

GFP-RAB5 transfected cells, grown on glass-bottom dishes (Ibidi), were imaged before and after a 20-minute treatment with Paprotrain (25 µM) using time-lapse microscopy. For Tf imaging, cells grown on glass-bottom dishes and treated with DMSO or Paprotrain for 15 min, were exposed to Tf-Cy3 (5 µg/mL) for 5 more minutes before imaging.

For the kinetics experiment of Paprotrain addition, GFP-RAB7 transfected cells, grown on glass-bottom dishes (Ibidi), were imaged every minute for 20 minutes following the addition of Paprotrain.

Time-lapse imaging was performed at 37°C using a spinning-disk microscope mounted on an inverted motorized microscope (Eclipse Ti-E, Nikon) through a 100 × 1.4NA PL-APO objective lens. The apparatus is composed of a Yogogawa CSU-22 spinning-disk head, a QuantEM EMCCD camera for image acquisition and Metamorph software (MDS) to control the setup.

#### Spreading assay

Huh-7 cells were treated for 2 h with 10 µM Paprotrain or the DMSO vehicle, then detached using Trypsin-EDTA and allowed to re-adhere on Matrigel-coated coverslips (20 µg/mL) in medium +/- Paprotrain. After 45 min, cells were fixed with PFA. Immunostaining was performed using anti-Phospho-Paxillin and -β1 Integrin antibodies to label focal adhesions. A sum projection of four optical slices at the plasma membrane-contacting surface was generated. Phospho-Paxillin staining was used to create a mask using Ilastik and the raw intensity of β1 Integrin within this mask was measured. Focal adhesion length was manually quantified based on Phospho-Paxillin labelling in a blinded manner. The adhesion surface was measured by outlining the perimeter of the cells. For large-scale quantification, an automated analysis was performed based on Phalloidin labelling using the software QuPath and CellPose.

Alternatively, cell adhesion was assessed by live-cell imaging, monitoring F-actin distribution during the spreading process. Huh-7 cells were treated for 2 h with 25 µM Paprotrain or the DMSO vehicle and SiR-Actin (1 µM, Spirochrome), a live cell actin probe, was added during the second hour before cells were detached and re-suspended on pre-coated Matrigel Ibidi chamber slides (20 μg/ml), in medium with 500 nM SiR-Actin +/- Paprotrain. The samples were allowed to settle for 30 min to establish multi-positions for the time-lapse. Sequences of images were taken for 1 h, 1 frame every 2 minutes with the inverted Confocal Zeiss LSM 980.

### Quantification and statistical analysis

#### Image processing

The images have been processed consistently and uniformly across data using ImageJ. Some processing, such as background subtraction and median filtering, was applied, while maintaining uniformity and preserving the integrity of the results.

#### Segmentation with pixel Classification workflow

Segmentation of fluorescence images was performed using the pixel classification workflow of Ilastik (version 1.4.1b6) to train a machine-learning-based model on manually annotated regions of interest. Intensity edge and texture features were extracted, and the trained classifier was applied to segment the entire dataset. Data was exported as simple segmentation and converted to binary mask in ImageJ

#### Co-localization between KIF20A and endosomal marker

Pearson coefficient quantification was performed using the BIOP JACoP plugin (Bolte and Cordelières, 2006) on the entire cell, with regions of interest (ROIs) defined in ImageJ.

#### Endosome area

To quantify the endosome area, a max intensity projection through the z-axis of 3 confocal images (corresponding to a total height of 0,42 µm) was done using the Image J software. Then segmentation was performed using Ilastik. Single vesicles were then segmented using the “Analyse Particles” plugin in ImageJ, with a minimum size threshold of 4 pixels per vesicle. For LAMP1 vesicles, the total area of LAMP1 staining was measured and divided by the number of vesicles observed visually, due to the clustering of the vesicles preventing segmentation.

#### EEA1-associated KIF20A intensity

To quantify EEA1-associated KIF20A intensity in HeLa cells following KIF20A silencing, a sum projection of three optical slices was generated and segmented using the EEA1 channel with Ilastik. The integrated density of KIF20A fluorescence was then measured within the mask.

To quantify EEA1-associated KIF20A-GFP intensity in MDCK-GFP-AID cells, three individual optical sections along the z-axis were segmented for EEA1, and the integrated density of KIF20A-GFP fluorescence was then measured within the mask.

#### Tf recycling

The total internal Tf fluorescence per cell was measured by first summing the optical sections of the entire cell and then quantifying the raw intensity. For each experiment, data were normalized to the highest fluorescence value at t0 for each condition (DMSO and PP). All datasets were then pooled and further normalized so that t0 was set to 100%, and expressed as a percentage of Tf remaining over time.

#### Tf and EGF colocalization

To quantify the colocalization between Tf and EGF, an object-based colocalization approach was used. Each channel was segmented using Ilastik. Colocalization was then assessed using these binary images by calculating the intersection area of the two signals as a percentage of the total area of the segmented Tf images in ImageJ.

#### Statistical analysis

All data were generated from cells pooled from at least three independent experiments represented as (n), unless mentioned, in corresponding legends. Statistical analyses were conducted using GraphPad Prism 8.0.1. A normality test was performed first, and the selected tests are detailed in the figure legends. No sample was excluded. Cells were randomly selected.

## AUTHOR CONTRIBUTIONS

A.K conceived and managed the study; A.K and J.So contributed to fund the project; J.Se contributed to the conception of and performed the experiments, collected the data and analysed them; S.L. and S.C. contributed to experiments. C.D., S.M. and J.So provided support with experimental set ups and results interpretation. J.Se and A.K wrote the initial manuscript with contribution from C.D., S.M and J.So.

## ACKNOWLEDGMENTS

We are grateful to B. Vitre and B. Delaval for sharing with us their MDCK KIF20A-GFP^AID^ cell line; to Arnaud Echard and Corinne Albigès-Rizo for generous gifts of reagents. We thank Romain Morichon from the Cytometrie and Imagerie Saint Antoine facility, France Lam, Chloé Chaumeton and Romain Naillon from the IBPS I2PS facility for assistance and technical support on confocal microscopy. We thank current and previous members of the lab for reviewing the article and for helpful discussions and most particularly Dr F. Praz for her scientific support and A. Užuotaitė for those first experiments. We greatly acknowledge the Nikon Imaging Center at Institut Curie-CNRS, PICT-IBiSA, member of the France-BioImaging national research infrastructure.

## FUNDING AND ADDITIONAL INFORMATION

J.S was funded by a PhD fellowship from <Ministère de l’Enseignement Supérieur et la Recherche= and by a 4^th^ year PhD fellowship (6 months) from the ARC foundation. A.K has received funding from GEFLUC R16170DD (2018). Confocal imaging acquisition using the Zeiss LSM 980 were performed at the IBPS Imaging Photonic Paris Seine (I2PS) facility. The IBPS Imaging facility is supported by Region-Île-de-France, Sorbonne-University and CNRS. Imaging acquisition using the Fluoview 3000 confocal microscope and the inverted IX83 microscope were performed at the CISA Cytometrie and Imagerie Saint-Antoine facility. The CISA Imaging facility received financial support from ITMO Cancer of Aviesan as part of the 2021-2030 Cancer Control Strategy (Inserm).

## SUPPLEMENTAL INFORMATION

**Supplementary Figure S1:**
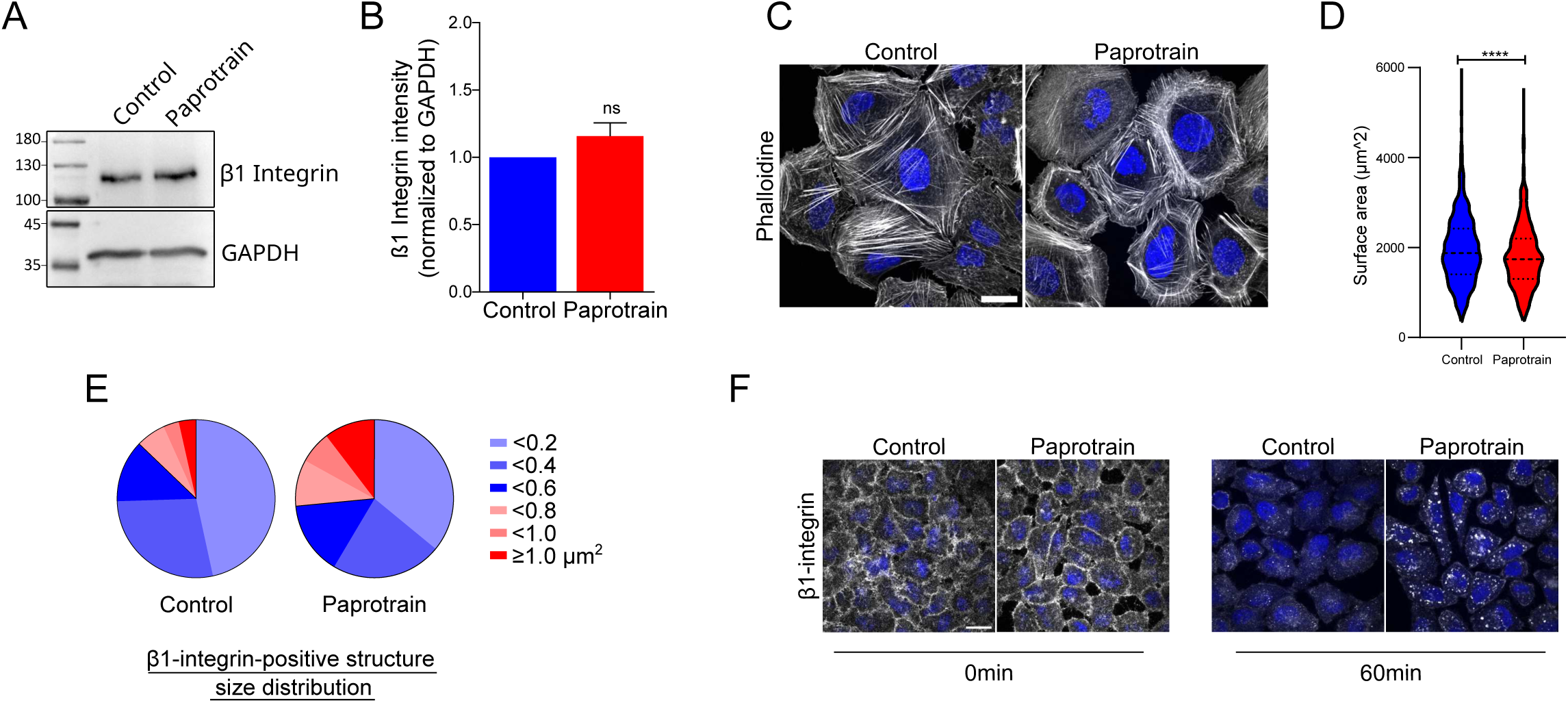
KIF20A activity is required for β1-Integrin trafficking, cell adhesion and migration. **(A-B)** Total β1-Integrin levels remain unchanged after KIF20A inhibition. Huh-7 cells were treated with DMSO (control) or Paprotrain (10 µM) for 2h, then detached and re -adhered onto Matrigel® (20 µg/mL) for 45 min. **(A)** Cell extracts were analyzed by Western blotting with β1-Integrin antibody. GAPDH is used as a loading control. **(B)** Quantification of **(A)** from three independent experiments. Data are presented as mean ± SEM. Unpaired t-test. ns = non-significant. **(C-D)** Huh-7 cells were treated with DMSO (control) or paprotrain (10 µM) for 2h, then detached and re-adhered onto Matrigel® (20µg/mL) for 45 min. Cells were fixed and stained with Phalloidin Alexa Fluor™ 633. **(D)** Quantification of the cell adhesion area. n= 500-1000 cells per condition from three independent experiments, mean ± SEM. Statistical analysis was performed using a Mann-Whitney test ****p<0.0001. **(E)** The distribution of β1- Integrin vesicles according to their surface area was analysed in both conditions (DMSO and Paprotrain) in Huh-7 cells across three independent experiments, n=58-60 cells. The data show the frequency of vesicles across different size ranges. **(F)** KIF20A inhibition induces intracellular accumulation of endocytosed β1-Integrin in HeLa cells. HeLa cells were serum-starved, treated with DMSO (control) or Paprotrain (25 µM) for 2h, and incubated with β1- Integrin-A488 for 45 min at 4°C. After washing, cells were fixed immediately (0 min) or after 1h at 37°C (60 min). Scale bar = 10 µm.

**Supplementary Figure S2:**
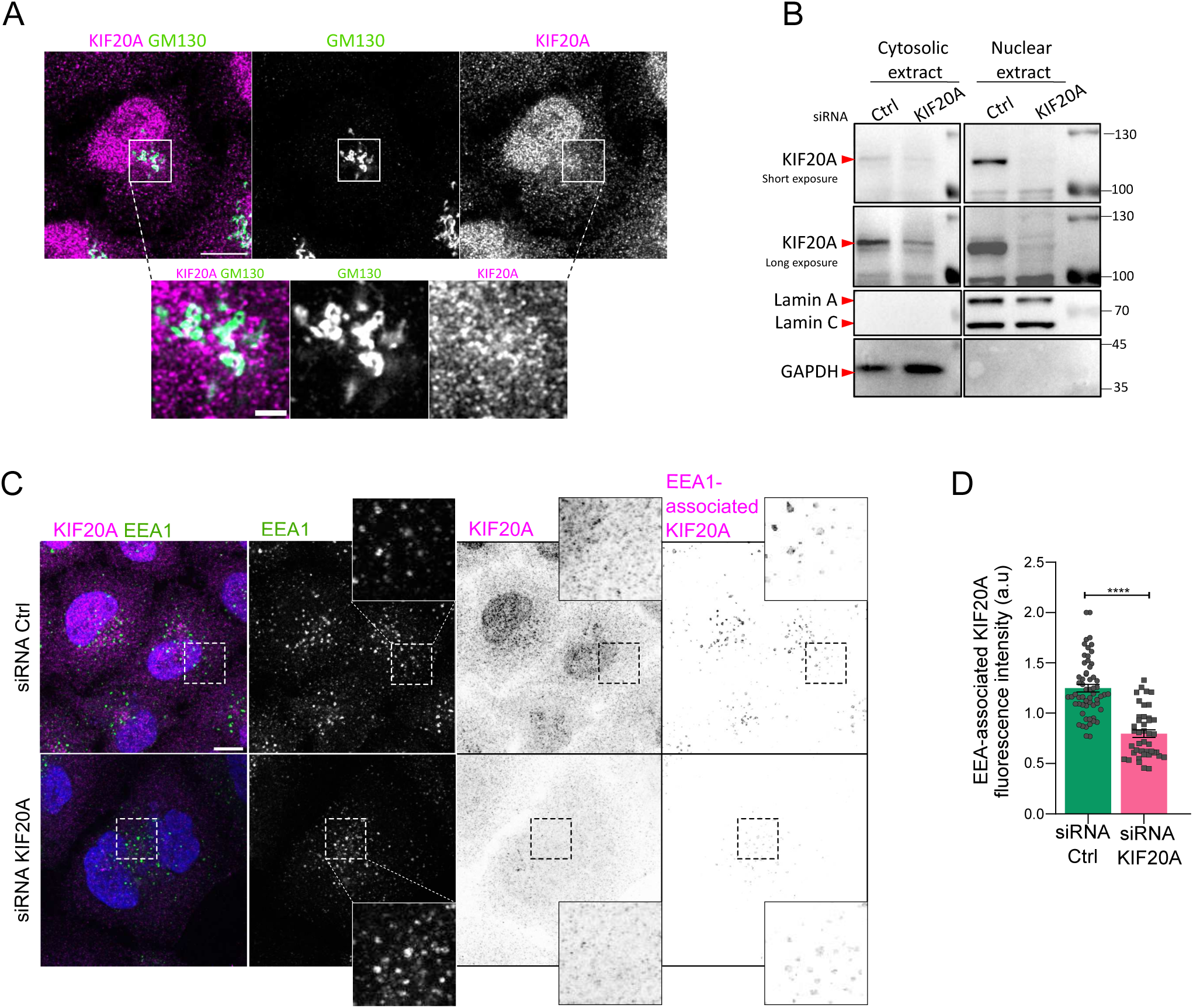
KIF20A localisation at endosomes. **(A)** HeLa cells were fixed with 4% PFA, and immunolabeled for endogenous KIF20A and GM130. Scale bar = 10 µm, magnified inset scale bar = 2 µm. **(B)** HeLa cells were treated with either control siRNA (Ctrl) or siRNA targeting KIF20A for 48h. Cellular fractionation was performed, separating cytosolic and nuclear fractions, prior to western blot analysis of KIF20A. Nuclear Lamins A/C and cytosolic GAPDH were used as controls. KIF20A is predominant in the nuclear fraction and efficiently knocked down by the siRNAs. **(C)** HeLa cells were treated as in **(B)**, fixed, and immunolabeled with antibodies against KIF20A and EEA1. As KIF20A is essential for cytokinesis, its depletion results in the formation of binucleated cells. The images represent a summation projection over 3 confocal slices. Scale bar = 10 µm. The EEA1- associated KIF20A was extracted from an EEA1 mask to show only the KIF20A associated with early endosomes. **(E)** Quantification of KIF20A fluorescence intensity on EEA1 mask. Data are represented as scatter plot and mean ± SEM. n=43-60 cells from two independent experiments. Statistical analysis was performed using a Mann-Whitney test, ****p<0.0001.

**Supplementary Figure S3:**
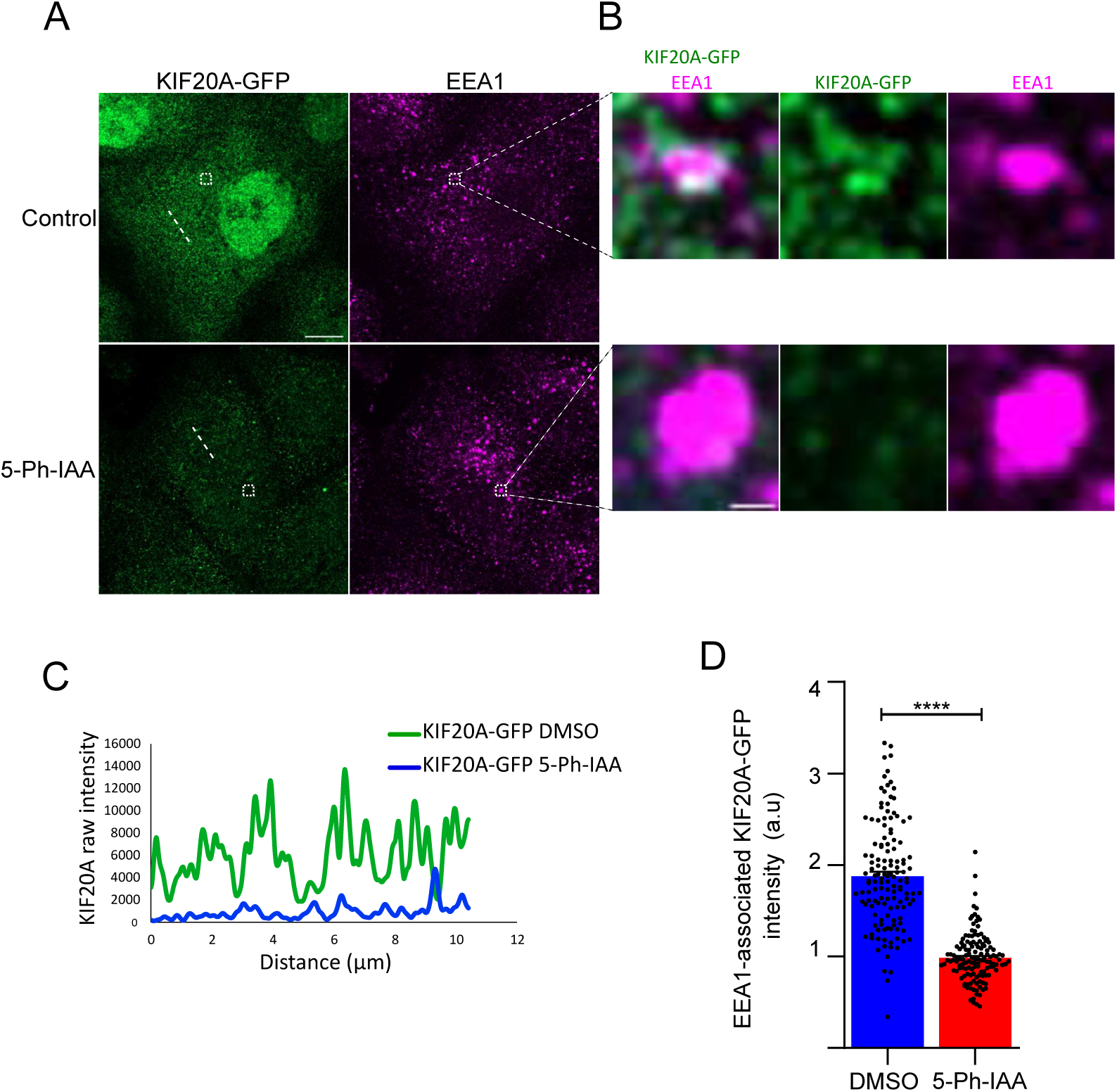
KIF20A activity is required for endosomal homeostasis. **(A-B)** MDCK KIF20A-GFP^AID^ were treated for 2h with DMSO (Control) or 5-Ph-IAA (auxin analog, 2 µM) to enable KIF20A degradation, and immunostained with antibodies against GFP and EEA1 **(A)**. Confocal microscopy images. Scale bar = 10 µm. The magnified view shows the colocalization between KIF20A and EEA1 **(B)**. **(C)** Line profiles (along white dash lines drawn in **A**) of GFP fluorescence intensity in control condition (green) and in 5-Ph-IAA treated cells (blue) showing the reduction of KIF20A-GFP^AID^ intensity, indicating degradation of KIF20A following 5-Ph-IAA induction. **(D)** Quantification of GFP intensity on an EEA1 mask. For each cell, three optical sections along the Z-axis were segmented for EEA1, and the intensity of GFP (KIF20A) was measured within the EEA1 mask (total intensity/EEA1 surface). Dot represents one slice. n=45 cells from 2 independent experiments. Data are represented as mean ± SEM. Statistical analysis was determined using a Mann-Whitney test ****p<0.0001.

**Movie S1: Dynamics of F-actin during cell adhesion in control condition**

Huh-7 cells were treated with DMSO for 2 hours. SiR-actin was used to label F-actin, and added on cells 1h prior to detachment. After trypsinisation, cells were allowed to adhere on Matrigel®- coated plates for 30 minutes and then imaged by time-lapse confocal microscopy. Frame rate: 1 image every 2 minutes, 9 planes with an increment of 0.5 µm z-projected per time point. Scale bar = 10 µm.

**Movie S2: Dynamics of F-actin during cell adhesion in KIF20A inhibited condition**

Huh-7 cells were treated with DMSO for 2 hours. SiR-actin was used to label F-actin, and added on cells 1h prior to detachment. After trypsinisation, cells were allowed to adhere on Matrigel®- coated plates for 30 minutes and then imaged by time-lapse confocal microscopy. Frame rate: 1 image every 2 minutes, 9 planes with an increment of 0.5 µm z-projected per time point. Scale bar = 10 µm.

**Movie S3: Dynamics of transferrin positive endosomes in control cells**

HeLa cells treated with DMSO were loaded with fluorescent Tf for 5 minutes and then imaged by time-lapse spinning disk confocal microscopy. Frame rate: 1 image per second, 3 planes with an increment of 0.3 µm z-projected per time point. Scale bar = 10 µm. The image displayed in Fig. 3 H is extracted from this movie.

**Movie S4: Dynamics of transferrin positive endosomes in KIF20A inhibited cells**

HeLa cells treated with Paprotrain were loaded with fluorescent Tf for 5 minutes and then imaged by time-lapse spinning disk confocal microscopy. Frame rate: 1 image per second, 3 planes with an increment of 0.3 µm z-projected per time point. Scale bar = 10 µm. The image displayed in Fig. 3 H is extracted from this movie.

**Movie S5: Dynamics of RAB5 positive endosomes in control cells**

HeLa cells transfected with GFP-RAB5 were imaged by time-lapse spinning disk confocal microscopy. Frame rate: 1 image every 2 second, 3 planes with an increment of 0.3 µm z-projected per time point. Scale bar = 10 µm. The image displayed in Fig. 3 I is extracted from this movie.

**Movie S6: Dynamics of RAB5 positive endosomes in KIF20A inhibited cells**

HeLa cells transfected with GFP-RAB5 were treated with Paprotrain for 20 minutes and then imaged by time-lapse spinning disk confocal microscopy. Frame rate: 1 image every 2 second, 3 planes with an increment of 0.3 µm z-projected per time point. Scale bar = 10 µm. The image displayed in Fig. 3 I is extracted from this movie.

